# Identification of COVID-19-relevant transcriptional regulatory networks and associated kinases as potential therapeutic targets

**DOI:** 10.1101/2020.12.23.424177

**Authors:** Chen Su, Simon Rousseau, Amin Emad

## Abstract

Identification of transcriptional regulatory mechanisms and signaling networks involved in the response of host to infection by SARS-CoV-2 is a powerful approach that provides a systems biology view of gene expression programs involved in COVID-19 and may enable identification of novel therapeutic targets and strategies to mitigate the impact of this disease. In this study, we combined a series of recently developed computational tools to identify transcriptional regulatory networks involved in the response of epithelial cells to infection by SARS-CoV-2, and particularly regulatory mechanisms that are specific to this virus. In addition, using network-guided analyses, we identified signaling pathways that are associated with these networks and kinases that may regulate them. The results identified classical antiviral response pathways including Interferon response factors (IRFs), interferons (IFNs), and JAK-STAT signaling as key elements upregulated by SARS-CoV-2 in comparison to mock-treated cells. In addition, comparing SARS-Cov-2 infection of airway epithelial cells to other respiratory viruses identified pathways associated with regulation of inflammation (MAPK14) and immunity (BTK, MBX) that may contribute to exacerbate organ damage linked with complications of COVID-19. The regulatory networks identified herein reflect a combination of experimentally validated hits and novel pathways supporting the computational pipeline to quickly narrow down promising avenue of investigations when facing an emerging and novel disease such as COVID-19.

## INTRODUCTION

Host responses to various insults is regulated by distinct sets of regulatory networks coordinating responses matched to the insult. Viral infections of human cells lead to the production of interferons (IFNs) as an antiviral mechanism [1]. TRIF, RIG-I and MDA-5-mediated activation of Interferon response factors (IRFs) responsible for the expression of antiviral genes, such as type I, II and III IFNs, are amongst critical regulators of antiviral immunity. In turn, Type I, II and III interferons will activate JAK-STAT signaling to further promote antiviral host responses [2]. This response must be kept in balance as viral clearance mechanisms can lead to tissue damage if not kept in check [3]. This balance can be especially hard to maintain in the presence of new emerging infections such as the novel coronavirus severe acute respiratory syndrome coronavirus - 2 (SARS-CoV-2) responsible for coronavirus disease 2019 (COVID-19), for which the host is naïve. The loss of a measured response can lead to severe complications of viral illnesses such as severe acute respiratory distress syndrome (ARDS), which has been observed in COVID-19 patients [1].

Unraveling the gene expression programs involved in the response of the host to the infection by SARS-CoV-2 can provide a fundamental understanding of COVID-19 and its complications and can enable identification of therapeutic targets and novel treatments. The transcriptional regulatory network (TRN), composed of transcription factors (TFs) and their target genes, play significant roles in regulating these gene expression programs. Computational reconstruction of ‘COVID-19-relevant’ TRNs that depict regulatory influence of TFs on genes differentially expressed due to infection of cells by SARS-CoV-2 would facilitate our understanding of this disease. While comparing the transcriptomic profiles of cells infected with SARS-CoV-2 with normal cells can provide some insight into the host response, understanding the specific response that leads to ARDS and other complications prominent in COVID-19 requires evaluating these molecular profiles against other respiratory viruses.

To identify transcriptional regulatory mechanisms involved in the host response to infection by SARS-CoV-2, we analyzed gene expression profiles of human lung epithelial cell lines that were mock-treated or infected by a respiratory virus (SARS-CoV-2, RSV, H1N1 and HPIV3) from a recent study [4]. SARS-CoV-2 (also named 2019 novel coronavirus (2019-nCoV) or human coronavirus 2019 (hCoV-19)) is a positive-sense single-stranded RNA virus, part of the broad coronavirus family. Similar to SARS-Cov-1 and Middle East respiratory syndrome (MERS), SARS-CoV-2 can cause severe acute respiratory disease in humans [1]. Respiratory syncytial virus (RSV) is a single-stranded negative-sense virus, a common cause of mostly mild respiratory disease in children. However, both in children [5] and adults [3], it can lead to serious lung diseases including ARDS. The influenza A virus H1N1, a negative-sense RNSA virus member of the orthomyxoviridae family, that was responsible for the 2009 swine flu pandemic. Human parainfluenza viruses (HPIV) are negative-sense RNA viruses that cause lower respiratory infections in children, chronically ill and elderly patients [6].

First, using a computational tool that we recently developed for reconstruction of ‘phenotype-relevant’ TRNs (InPheRNo) [7], we reconstructed COVID-19-relevant TRNs and identified key regulatory TFs involved in the progress of the disease. TRNs are network representations of regulatory mechanisms in a cell, in which nodes are TFs or genes, and each TF-gene edge represents a regulatory effect of the TF on the gene. Unlike other methods for reconstruction of TRNs that are usually agnostic to the phenotype under investigation, InPheRNo utilizes probabilistic graphical models to directly incorporate phenotypic information in the TRN reconstruction. This approach enables identification of transcriptional regulatory mechanisms that are involved in the specific phenotype under investigation by expression profiling (in this study, response of cells infected by SARS-CoV-2 as compared to mock-treated cells or cells infected by other viruses). Our results identified known and novel key regulatory TFs and signaling pathways involved in COVID-19 and its associated complications.

Next, using a network-guided approach based on random walks on graphs, we identified kinases that are most associated with the reconstructed COVID-19-relevant TRNs, as regulators of these networks and potential therapeutic targets. Kinases are enzymes that are involved in the regulation of protein activities through phosphorylation and are a major category of drug targets for human diseases. Using data from gene knockdown experiments from the LINCS database [8], we observed that these kinases indeed influence the expression of genes in the reconstructed TRNs in epithelial cells. Our analyses using network-based algorithms and machine learning tools provided a systems biology perspective of the response of the epithelial cells to infection by SARA-CoV-2 and identified regulatory mechanisms specific to this virus. In addition, our results implicated important families of kinases (including JAK and MAPK family) that may be used as therapeutic targets for COVID-19.

## RESULTS

### COVID-19-relevant transcriptional regulatory networks implicate major transcription factors involved in SARS-CoV-2 host response to infection

We sought to identify transcriptional regulatory mechanisms involved in the host response to SARS-CoV-2 infection. For this purpose, we obtained gene expression profiles of human lung epithelial cells that were mock-treated or infected by SARS-CoV-2, respiratory syncytial virus (RSV), human parainfluenza virus type 3 (HPIV3), influenza A/Puerto Rico/8/1934 (H1N1) virus (IAV), and IAV that lacks the NS1 protein (IAVdNS1) from a recent study [4]. The epithelial cells were Normal Human Bronchial Epithelial (NHBE), transformed lung alveolar (A549), A549 cells transduced with a vector expressing human ACE2 (A549-ACE2), and Calu3 cells.

To identify COVID-19-relevant TRNs, we used InPheRNo [7], a method that we recently developed to identify ‘phenotype-relevant’ TRNs using gene expression profiles of multiple samples and their phenotypic labels. InPheRNo is based on a probabilistic graphical model (PGM) designed to integrate the collective regulatory influence of multiple TFs on a gene with the association of the gene’s expression with a phenotype to identify regulatory mechanisms that are phenotype-relevant (as opposed to phenotype-independent). In this approach, first the p-values of gene-phenotype associations (e.g. using differential expression analysis) and p-values of gene-TF associations (using a two-step procedure based on the Elastic Net algorithm) are obtained and provided as input ‘observed variables’ to the PGM. The PGM is then trained on the data to obtain posterior probabilities for each TF-gene pair determining whether the TF regulates the gene in a phenotype-relevant manner (details are provided in the original manuscript [7]).

Using data corresponding to SARS-CoV-2 infected samples and their corresponding mock-treated control, we first performed differential expression analysis and then reconstructed a COVID-19-relevant TRN associated with SARS-CoV-2 infection (as compared to mock-treated samples) using 500 most differentially expressed genes (FDR < 1.42E-3, shown in Supplementary Table S1), henceforth called ‘SvM’ (SARS-CoV-2 versus mock-treated). To also identify regulatory mechanisms that are specific to SARS-CoV-2 infection (as opposed to infection by other viruses), we used InPheRNo to reconstruct a COVID-19-relevant TRN using 500 differentially expressed genes (FDR < 1.43E-3, Supplementary Table S1) by analyzing data corresponding to epithelial cells infected by SARS-CoV-2, IAV, IAVdNS1, RSV, and HPIV3. Henceforth, we use ‘SvOV’ (SARS-CoV-2 versus other viruses) to refer to the second network. The details of the analysis are provided in Methods and the reconstructed networks are provided in Supplementary Table S2.

Given the SvM and SvOV COVID-19-relevant TRNs, we ranked TFs based on the number of COVID-19-relevant target genes within each network. Tables 1-2 show the ranked list of top 8 TFs and top 21 TFs that target at least 1% of the considered COVID-19-related genes in SvM and SvOV networks, respectively (see Supplementary Table S3 for the full ranked list). We compared the targets of these TFs identified by InPheRNo, with their targets determined using ChIP-seq data available in the Gene Transcription Regulation Database (GTRD) database [9]. Three of the top 8 TFs in the SvM network (NFKB1, STAT1, RELB) were present in the GTRD dataset. Out of the 18 targets identified by InPheRNo for these TFs, 17 were confirmed by GTRD (p= 8.62E-07, hypergeometric test). Similarly, four of the top TFs in the SvOV network (STAT1, RCOR1, EGR1, ZNF512B) were present in this dataset. Out of the 45 targets found for these TFs by InPheRNo, 37 were confirmed using GTRD (p = 2.36E-15, hypergeometric test).

**Table 1:** Top 8 TFs implicated in the SvM (SARS-CoV-2 vs. mock-treated) network. The TFs are ranked based on the number of their COVID-19-relevant target genes identified by InPheRNo. The second column shows the percent of the considered genes that each TF regulates.

| Transcription Factors | Percent of target genes |
| --- | --- |
| IRF9 | 3.05% |
| IRF7 | 1.96% |
| NFKB1 | 1.74% |
| MAFF | 1.53% |
| SP110 | 1.09% |
| RELB | 1.09% |
| STAT1 | 1.09% |
| BATF2 | 1.09% |

**Table 2:** Top 21 TFs implicated in the SvOV (SARS-CoV-2 vs. other viruses) network. The TFs are ranked based on the number of their COVID-19-relevant target genes identified by InPheRNo. The second column shows the percent of the considered genes that each TF regulates.

| Transcription Factors | Percent of target genes |
| --- | --- |
| STAT1 | 5.89% |
| STAT2 | 2.95% |
| MLX | 2.74% |
| EGR4 | 1.47% |
| RCOR1 | 1.47% |
| SP140L | 1.47% |
| TP53 | 1.26% |
| RCOR2 | 1.26% |
| MAX | 1.26% |
| ZNF496 | 1.26% |
| ZNF512B | 1.05% |
| SMAD7 | 1.05% |
| SOX12 | 1.05% |
| IRF2 | 1.05% |
| HDX | 1.05% |
| EGR1 | 1.05% |
| SP110 | 1.05% |
| IRF9 | 1.05% |
| ZNF143 | 1.05% |
| NFIX | 1.05% |
| ZBTB32 | 1.05% |

Encouragingly, many of these TFs identified by InPheRNo have been previously shown to be activated during COVID-19 or infections by other viruses. For example, Interferon regulatory factor 9 (IRF9), the top hit in Table 1, was shown to be activated in SARS-CoV-2 infected NHBE cells [10]. While interestingly, in contrast to observations with SARS-CoV-1, infection by SARS-Cov-2 failed to limit STAT1 phosphorylation [11], suggesting that STAT1 activity is maintained in SARS-CoV-2 CaLu-3 infected cells.

### Functional characterization of COVID-19-relevant TRNs implicate major signaling pathways involved in the disease

In order to determine the functional characteristics of gene expression programs involved in COVID-19, we performed pathway enrichment analysis for the implicated TFs and their COVID-19-relevant target genes in the SvM and SvOV networks. For this purpose, we used the gene set characterization (GSC) computational pipeline of KnowEnG (Knowledge Engine for Genomics) analytical platform [12]. The GSC pipeline enables ‘standard’ gene-set enrichment analysis (using Fisher’s exact test), as well as advanced ‘network-guided’ analysis. The network-guided mode is an implementation of an algorithm called DRaWR [13], which utilizes random walk with restarts (RWR) algorithm on a user-selected gene interaction network to rank pathways based on their relevance to a query gene-set. Including a gene interaction network (e.g. a protein-protein interaction (PPI) network) enriches the analysis and enables identification of important pathways that may not be detectable using simple overlap-based Fisher’s exact test.

To this end, we used the set of top TFs in the SvM and SvOV networks (Tables 1-2) as separate query sets and performed standard and network-guided (using experimentally verified PPI network from STRING database [14]) pathway enrichment analysis (using Reactome pathways [15]) with default parameters (Supplementary Table S4). Pathways related to cytokine signaling and interferon signaling (interferon gamma signaling and interferon alpha/beta signaling) were implicated for both TRNs and using standard and network-guided analysis. Next, we repeated the network-guided analysis above by considering each TF and the set of its target genes in the SvM and SvOV networks as a separate query gene set (Fig. 1 and Supplementary Tables S5-S6). Fig. 1 shows gene sets that are implicated for at least two TFs and their targets in each TRN. Pathways related to Immune system, cytokines and interferon signaling were again among the pathways implicated for the majority of TFs (and their COVID-19-relevant targets).

**Figure 1:**
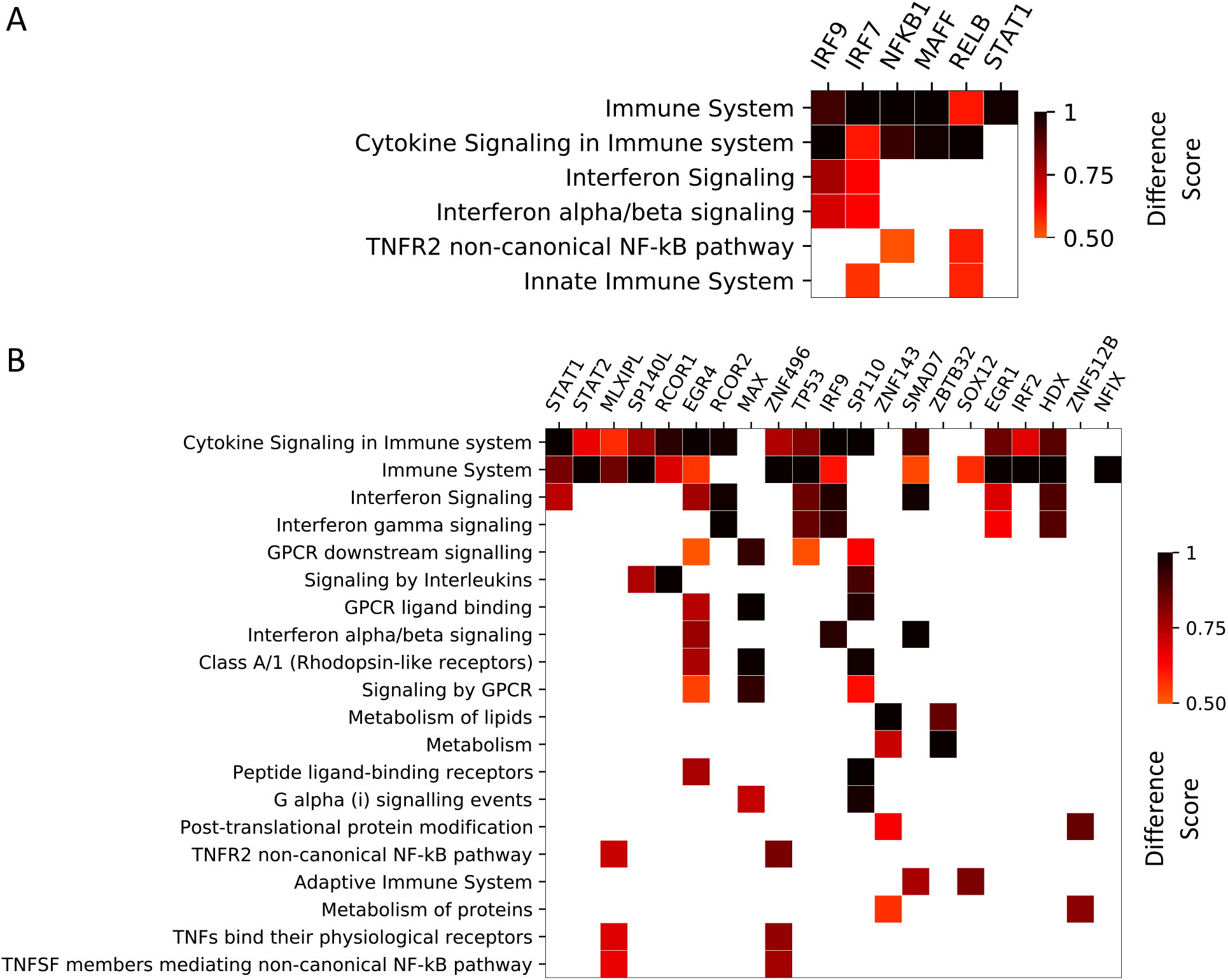
Pathway enrichment analysis using network-guided gene set characterization pipeline of KnowEnG. The columns correspond to top TFs and their COVID-19-relevant targets identified using InPheRNo. Only pathways that have been implicated for at least two TFs (and their targets) are depicted (see Supplementary Tables S5 and S6 for the full list). The heatmap shows the ‘difference score’ assigned to each pathway using KnowEnG. Cases in which the score was <0.5 are shown as white. A) Implicated pathways for top TFs in the SvM TRN. B) Implicated pathways for top TFs in the SvOV network.

### Identification of kinases associated with COVID-19-relevant TRNs as potential therapeutic targets

Kinases are enzymes that are involved in the regulation of protein activities through phosphorylation and are a major category of drug targets for human diseases [16]. Consequently, we sought to identify human kinases that are most associated with the constructed COVID-19-relevant TRNs as important signal transducers for this disease. For this purpose, we formed a kinase-substrate interaction (KSI) network by aggregating kinase-substrate relationships from three previous studies [17–19] (see Methods for details). The aggregated KSI contained 29594 kinase-substrate relationships corresponding to 406 unique kinases and 3942 unique substrates.

To identify kinases most associated with COVID-19 we used foRWaRD, a computational tool that we recently developed to rank nodes and sets of nodes in a heterogenous network based on their relevance to a set of query set using random walk with restarts (RWR) [20]. As input to foRWaRD, we provided the aggregated KSI, a gene-gene interaction network (here we used HumanNet integrated network [21]), and a query set containing the top TFs obtained from SvM TRN (8 TFs) and SvOV TRN (21 TFs), separately. foRWaRD first forms a heterogenous network by superimposing the substrates on their corresponding gene nodes in the gene-gene interaction network. Then, it performs two runs of the RWR algorithm on this heterogenous network: one run with the query set (set of TFs) as the restart set and another run with all nodes as the restart set (to be used as control). Each run of the RWR provides a probability score for each node (including those corresponding to kinases), representing the relevance of the node to the restart set. Finally, a normalized score for each kinase is obtained by comparing the scores of the two runs of the RWR, and kinases are ranked based on how much their query set score is higher than their background (i.e. control) score. Table 3 shows the 15 highest ranked kinases for the top TFs corresponding to the SvM network and top TFs corresponding to the SvOV network, respectively. The full ranked lists of kinases are provided in Supplementary Table S7. Figs. 2 and 3 show network representations of the interactions among these kinases, their substrates, and the COVID-19-relevant TRNs. Fig. 2 only includes direct kinase-TF interactions, while Fig. 3 and Supplementary Table S8 include indirect interactions of kinases and TFs.

**Table 3:**
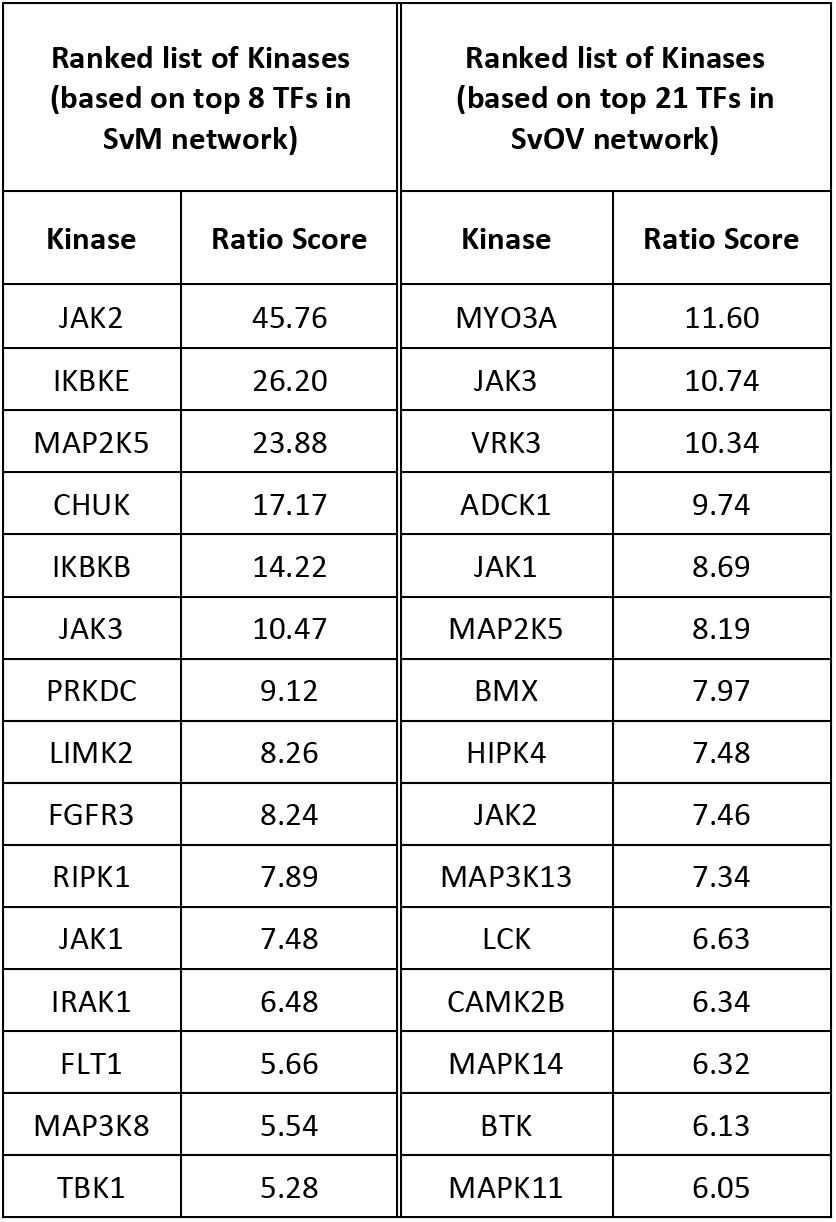
Top 15 kinases identified using foRWaRD for the top TFs in SvM and SvOV networks. The kinases are ranked based on the ratio of their query set probability score to their background probability score. Any ratio score >1 implies that the kinase is scored higher using the top TF query set compared to its control.

**Figure 2:**
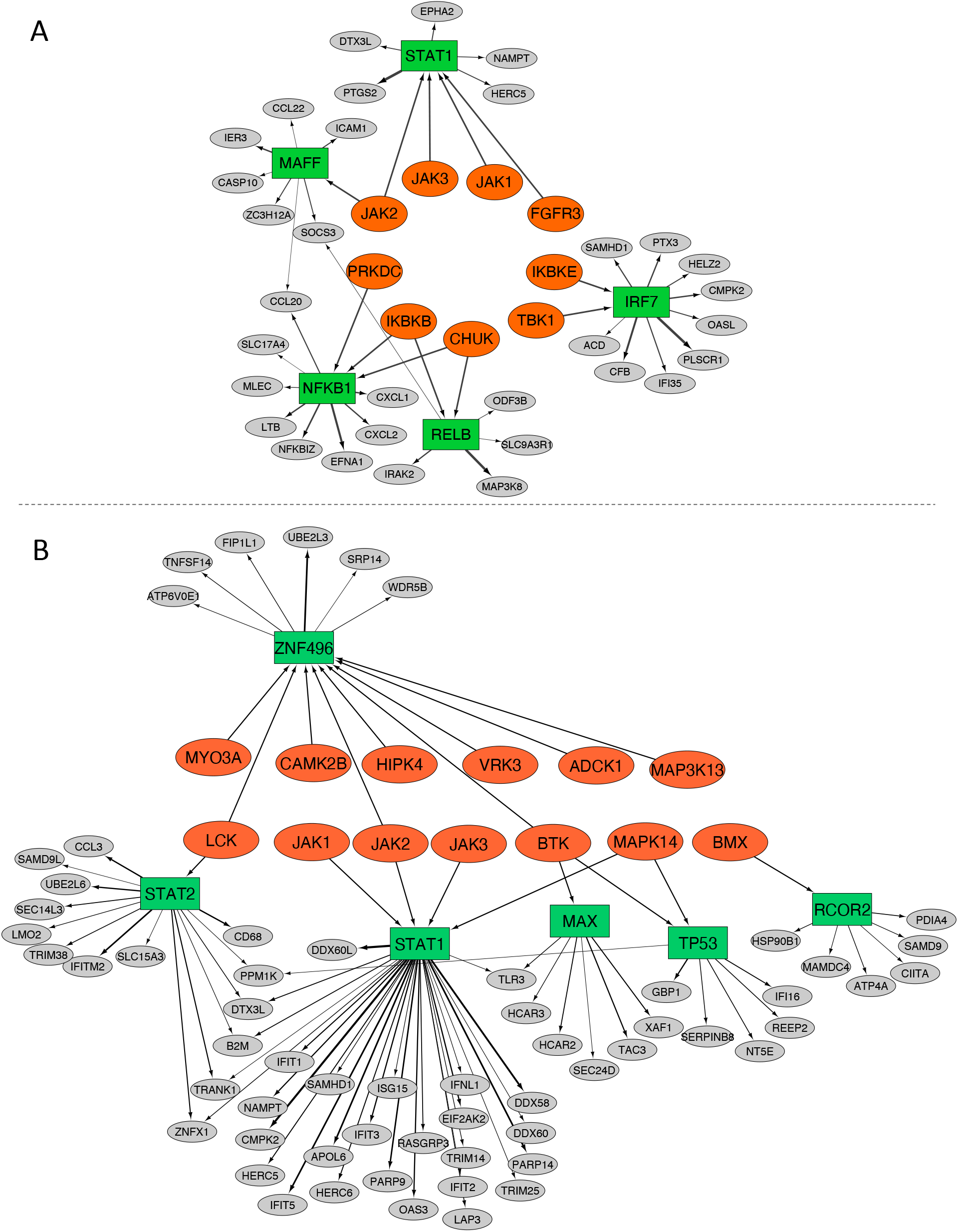
Network representation of direct interactions between implicated kinases, TFs and genes associated with COVID-19. Kinases are depicted as orange ellipses, TFs are depicted as green rectangles and target genes are depicted as grey ellipses. Only direct kinase-TF interactions present in the aggregated KSI are depicted. A) Network corresponding to the SvM analysis. B) Network corresponding to the SvOV analysis.

**Figure 3:**
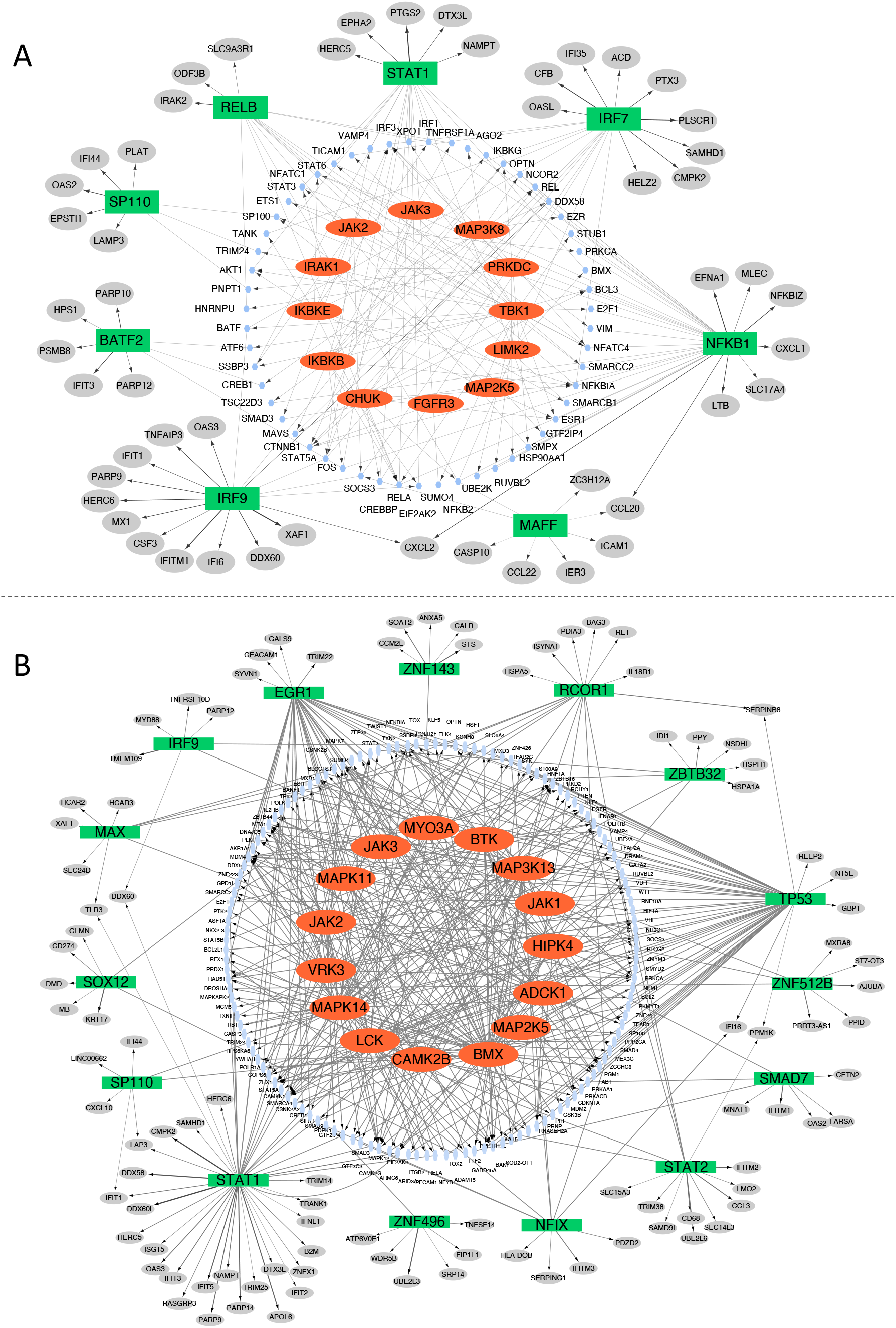
Network representation of indirect interactions between implicated kinases, substrates, TFs and genes associated with COVID-19. Kinases are depicted as orange ellipses, substrates that interact with at least one of the implicated TFs in the HumanNet integrated network are depicted as blue ellipses, TFs are depicted as green rectangles and target genes are depicted as grey ellipses. Directed edges show interactions between kinases and their substrates (obtained from the aggregated KSI) as well as TFs and their target genes (obtained using InPheRNo). Undirected edges correspond to interactions between substrates and TFs (obtained from HumanNet Integrated network). A) Network corresponding to the SvM analysis. B) Network corresponding to the SvOV analysis.

As can be seen in Table 3, several families of kinases are implicated in both networks. JAK1, JAK2, and JAK3, which are identified among the top 15 kinases for both SvM and SvOV TRNs, belong to the Janus kinase family, a family of non-receptor tyrosine kinases [22]. This family of kinases are involved in the transduction of cytokine-mediated signals through the JAK-STAT pathway. The members of the Janus family and the JAK-STAT pathway have been suggested as potential therapeutic targets in COVID-19 [23–26], supporting the validity of the results from this analysis.

### Evaluation of the predicted kinase-gene relationships using gene knockdown experiments

Since foRWaRD incorporates both direct and indirect interactions to identify kinases, we sought to determine whether the knockdown of identified kinases directly influence the expression of the TFs and their target genes in the COVID-19-relevant TRNs. To this end, we obtained gene expression signatures corresponding to shRNA knockdown experiments from the LINCS dataset [8]. We only focused on experiments performed in A549 cell line, since it is one of the cell lines used in our analysis to construct the COVID-19-relevant TRNs and a cell line shown to be susceptible to SARS-CoV-2 infection [27]. The gene expression signatures correspond to z-score normalized changes in the expression of 978 ‘L1000 landmark genes’ as a result of knockdown of a single gene, when compared to control (no knockdown). We defined a landmark gene to be positively (or negatively) influenced by the knockdown of a kinase if its expression increased (decreased) as a result of the knockdown and also if its normalized expression change was among the top (bottom) 15% of all landmark genes.

Out of the 15 kinases implicated using foRWaRD for the SvM analysis, shRNA knockdown signatures in A549 cells were available for 13 of them. In addition, 10 target genes and 2 TFs from the SvM TRN were among the L1000 landmark genes whose expression change were measured. All of the 2 TFs and the 10 target genes were positively or negatively influenced by the knockdown of at least one of the implicated kinases (Supplementary Table S9 and Supplementary Figures S1-S2), supporting the interactions discovered in this study.

We repeated the analysis above using the kinases implicated for the SvOV network. Knockdown signatures were only available for 7 (out of 15) implicated kinases. In addition, 14 target genes and 3 TFs from the SvOV TRN were among the L1000 landmark genes. Our analysis showed that 12 target genes (out of 14) and 2 TFs (out of 3) were positively or negatively influenced by the knockdown of at least one of the 7 kinases (Figure 4, Supplementary Table S9 and Supplementary Figure S3). Figure 4 shows the histogram of the expression changes of the landmark genes as a result of each kinase knockdown and its effect on the TFs and target genes in the SvOV TRN.

**Figure 4:**
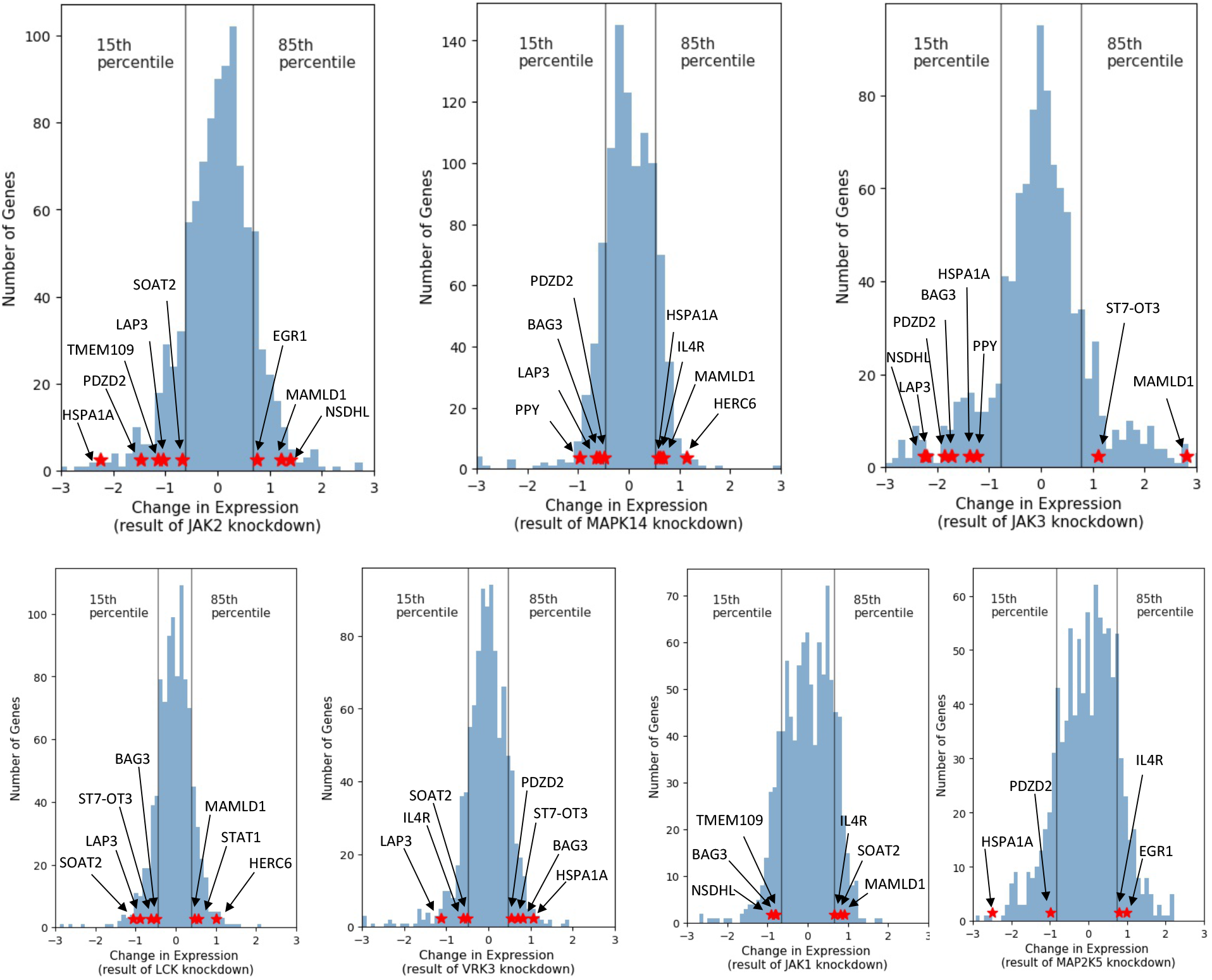
The histogram of z-score normalized gene expression changes of LINCS L1000 landmark genes in A549 cells as a result of knockdown of kinases implicated in the SvOV analysis. Each histogram corresponds to the knockdown of one kinase. Vertical lines depict the 15^th^ and 85^th^ percentiles and red stars show the TFs and target genes in the SvOV network that are positively or negatively influenced by the experiment.

Taken together, these results show that our computational pipeline that strings together InPheRNo, KnowEnG’s GSC, and foRWaRD is capable of identifying biologically plausible signaling networks involved in regulating the responsible of airway epithelial cells to SARS-CoV-2.

## DISCUSSION and CONCLUSION

The aim of this study was to identify signaling and transcriptional regulatory networks that play key roles in transducing signals specific to SARS-CoV-2 infection of airway epithelial cells in order to better understand the pathophysiology of COVID-19 and provide a list of potential molecular targets for therapies aimed at altering the clinical courses of the disease. The response was studied in comparison to mock-treated cells and other viruses (RSV, H1N1, HPIV3).

### Insights learned from the regulatory networks associated with SARS-CoV-2 versus mock-treated cells

In the first instance, comparing to mock-treated cells, we can identify the key regulatory pathways of antiviral responses. Alterations of these pathways (whether genetic, epigenetic or environmental) can have important consequences for a broad range of infections. If their activity is diminished or impaired, susceptibility to viral infections is expected, with high titers of virus likely increasing infectivity. This is the case for the loss of function of TLR7 [28] or type I and type III IFN-related genes [29, 30] leading to more severe COVID-19 disease in younger individuals. As expected, pathway enrichment analysis showed that immune (cytokine) signaling related to interferon were the top hits, as is expected for viral infection of host cells [31]. Amongst the lists of TFs identified is IRF9, a TF shown to be activated by SARS-CoV-2 [10] that forms a trimeric complex with STAT1 and STAT2 termed IFN-stimulated gene factor 3 (ISG3) that binds to the IFN stimulated response element (ISRE) and controls the expression of type I and III IFN essential for the control of Influenza A virus replication [32]. Interferons alpha and beta (IFN-a and IFN-b) are among the type I IFNs that regulate the activity of immune system and act as antiviral cytokines. A recent study has shown an association between impaired interferon type I response (represented by low activity of IFN-a and IFN-b) and severity of COVID-19 and has suggested blood deficiency in type I IFN as a hallmark of severe manifestation of the disease [28]. In addition, several studies have suggested type I IFNs as potential antiviral treatments for COVID-19 [29, 30]. IFN-g is a cytokine involved in innate and adaptive immunity and is the only member of type II IFNs. A recent study has shown a correlation between COVID-19 severity and a decrease in the production of IFN-g by CD4+ T cells [31].

Pathway enrichment provides a high-level view, which lacks granular information about the regulatory pathways involved. For this reason, we focused our analysis on linking protein kinases to the identified network using foRWaRD. This analysis confirmed the importance of JAK-STAT and NFkB signaling for the expression of IFN genes and antiviral responses in SARS-CoV-2 versus mock-treated cells (Fig. 2A and 3A). Identified kinases also included TANK-binding kinase 1 (TBK1) and IKKe, critical regulators of the transcription factor IRF7 that regulate amongst other genes IFN-induced oligodenylate synthetase-like (OASL) and Pentraxin-3 (PTX3) (Fig. 2A). TBK1/IKKe acts downstream of TLR3 and TLR7 via the TRIF-signaling adapter, two TLR identified in genetic studies to confer susceptibility to COVID-19 [33, 34]. Moreover, high circulating expression of PTX3, derived from monocytic and endothelial cells is a predictor of short-term COVID-19 mortality [35]. The analysis thus identified several relevant pathways to SARS-CoV-2 infection and disease susceptibility/severity.

### Insights learned from the regulatory networks associated with SARS-CoV-2 versus other respiratory virus-treated cells

Investigation of the regulatory networks specifically associated with SARS-CoV-2 can help better understand either acute complications or long-term consequences of COVID-19. This is particularly important for a novel disease, for which limited information is available. Considering that as of December 2020, more than 75 million people have been infected, the potential of diverging responses is huge and can have long-lasting impact on the health of many. Therefore, better understanding the peculiarities of SARS-CoV-2 compared to other viruses is paramount to deal with the coming fall out of the pandemic. When looking at differences between mock- and other viruses-treated cells, the enrichment analysis highlighted a potential role for GPCR signaling associated with the TFs MAX, SP110 and EGR4. GPCRs form a large family of 7-transmembrance cell surface receptors that mediates numerous cellular functions, including cytokine/chemokine signaling involved in leukocyte chemotaxis [36]. The association with the above-mentioned TFs postulate a role in controlling cell differentiation/cell death programs by GPCR.

In the SvOV network, additional protein kinases were identified including MAPK11, MAPK14, BTK, and BMX. MAPK11 and MAPK14 both belong to the p38 MAPK family which are involved in the cellular responses to extracellular stimuli including proinflammatory cytokines such as IL-6 [37]. Previously, it has been shown that one of the proteins expressed by SARS-CoV virus upregulates p38 MAPK [38]. A global phosphorylation analysis of SARS-CoV-2 infected epithelial cells also identified MAPK14 and MAPK11 as kinases upregulated during infection that make important contributions to host responses [39]. Accordingly, inhibition of MAPK14 and MAPK11 family has been proposed as a potential therapeutic approach in COVID-19 [37]. In addition to be a downstream target of TLR-signaling pathways including TLR3 in airway epithelial cells [40], MAPK14 regulates IL-6 expression and mRNA stability [41]. IL-6 circulating levels are elevated in severe COVID-19 [42]. Moreover, MAPK14 is an important signal transducer of IL-17 in endothelial cells [43], involved in neutrophilic inflammation, that may be important mediators of thrombosis in COVID-19 via the release of Neutrophil-Extracellular-Traps [20, 44]. MAPK14 is an important target of the potent corticosteroid dexamethasone [45], that was shown to decrease mortality in severe COVID-19 [46].

BMX and BTK are non-receptor tyrosine kinases that belong to the Tec kinase family. Tec family has been shown to be involved in the intracellular signaling mechanisms of cytokine receptors and antigen receptor signaling in lymphocytes [47]. BMX has been shown to link both MYD88, another TLR-signaling adapter, and Focal Adhesion Kinases, a kinase associated with integrin activation to the synthesis of IL-6 [48]. The inhibition of genes in this family, and particularly BTK [41], has been proposed as a therapeutic approach to protect COVID-19 patients against pulmonary injury [42] and to block thrombo-inflammation [43]. Two potential downstream targets of BTK according to our regulatory networks (Fig. 2B), Tumor necrosis factor ligand superfamily member 14 (TNFS14) and XIAP-associated factor 1 (XAF1) have been identified in a single cell transcriptomic study comparing IAV and SARS-CoV-2 responses [49]. TNFSF14 is a ligand of the lymphotoxin beta receptor that amplifies NFkB signaling in T lymphocytes to increase IFN-gamma production [50]. XAF1, as its name implies, binds XIAP (BIRC4) an important regulator of inflammatory signaling and apoptosis, that increases TRAIL-mediated apoptosis in response to IFNb [51]. While these responses are likely desirable during early phases of the infection, whether they can also contribute to immunopathology in the second sustained phase of the disease warrants further investigation.

In conclusion, our results obtained by stringing together three powerful computational tools (InPheRNo, KnowEnG GSC, and foRWaRD) identified regulatory networks, pathways, and kinases, many of which have already been associated with COVID-19 in previous studies. These results also provided further information on putative regulatory mechanisms underpinning the infection of epithelial cells by SARS-CoV-2 and identified novel potential therapeutic targets that can serve as the basis for future identification and development of drugs that mitigate the impact of COVID-19 in individuals at risk of severe complications. The SvM regulatory network was mostly related to classic antiviral response pathways (IRF, IFN, JAK-STAT, etc.) that are well supported by genomic data on diseases susceptibility to COVID-19 [33, 34]. Interestingly, the SvOV regulatory network identified pathways associated with regulation of inflammation (MAPK14) and immunity (BTK, BMX) that may contribute to exacerbate organ damage.

## METHODS

### Data collection

We downloaded mock-treated and infected RNA-seq gene expression profiles of human lung epithelial cells from the Gene Expression Omnibus (GEO) database (accession number: GSE147507). For the reconstruction of SARS-CoV-2 versus mock (SvM) TRN, we used 24 samples corresponding to independent biological triplicates of SARS-CoV-2 infected NHBE, A549, A549-ACE2, and Calu3 and their corresponding mock-treated control. For the reconstruction of SARS-CoV-2 versus other viruses (SvOV) TRN, we used 33 samples corresponding to independent biological replicates of A549 cells infected with SARS-CoV-2, RSV, IAV, and HPIV3, NHBE cells infected with SARS-CoV-2, IAV, and IAVdNS, A549-ACE2 cells infected with SARS-CoV-2, and Calu3 cells infected with SARS-CoV-2.

We downloaded the list of human TFs from AnimalTFDB [52]. Experimentally verified protein-protein interaction network from the STRING database [14] and HumanNet integrated network [21] were downloaded from KnowEnG’s knowledge network (version 17.06) available at the address https://github.com/KnowEnG/KN_Fetcher/blob/master/Contents.md. The list of target genes for the top TFs (identified using ChIP-seq) was downloaded from the GTRD database (http://gtrd.biouml.org/downloads/20.06/intervals/target_genes/Homo%20sapiens/genes%20promoter%5b-1000,+100%5d/). In this dataset, a gene is considered to be target of a TF if its promoter region (defined as the interval [-1000, +100] bp relative to gene’s transcriptional start site) contains at least one GTRD meta-cluster for the TF. The meta-clusters reflect ChIP peaks for the same TF-gene but integrated from different experiment conditions and different peak calling methods [9].

To form the aggregated KSI network, we obtained kinase-substrate relationships from three previous studies. The interactions corresponding to homo sapiens were downloaded from PhosphoSitePlus database (www.phosphosite.org) [17], PhosphoNetworks (www.phosphonetworks.org) [18], and the supplementary material of an independent study [19]. After removing duplicate edges, we formed a KSI involving 29594 kinase-substrate relationships corresponding to 406 unique kinases and 3942 unique substrates. LINCS Level 5 consensus signatures (‘trt_sh.cgs’) corresponding to shRNA knockdowns in A549 cell line were obtained from GEO with the accession number (GSE92742). We used the Consensus Gene Signatures (CGS) data since they correspond to gene expression changes that are common among multiple shRNAs that target the same gene, mitigating off-target effects [53].

### Reconstruction of COVID-19-relevant TRNs using InPheRNo

InPheRNo [7] is a computational method that utilizes a probabilistic graphical model to combine information on the significance (pseudo p-value) of gene-TF associations (from their expression profiles) with information on the significance (p-value) of gene-phenotype associations to construct phenotype-relevant TRNs. As input, it accepts a list of TFs, the expression profiles of genes and TFs (in multiple samples), and the p-value of association between genes’ expression and a phenotype.

To construct the TRNs using InPheRNo, we first performed differential expression analysis using EdgeR [54] with the cell type as a confounding factor. In the case of SvOV, since for different viruses the measurements were obtained at different time-points, we also included the time of measurement post infection as a confounding factor. Next, we quantile normalized the gene expression profiles using voom [55] and then z-score normalized the results. We constructed the SvM and SvOV TRNs using 500 most differentially expressed genes. We ran InPheRNo (downloaded from https://github.com/KnowEnG/InPheRNo) with 1000 iterations, 500 repeats and default values for other parameters.

### Pathway enrichment analysis

We used KnowEnG’s gene set characterization (GSC) pipeline [12] (www.knoweng.org/analyze) to perform pathway enrichment analysis. For the standard mode of GSC pipeline (without use of any gene interaction network), we chose Reactome pathways [15] as the option for target gene sets and the rest of parameters were left as default. We excluded pathways with smaller than 10 genes and adjusted the enrichment p-values (Fisher’s exact test) for multiple tests using Benjamini-Hochberg false discovery rate (FDR). Pathways with FDR < 0.05 were considered statistically significant.

For the knowledge-guided mode of the GSC pipeline, we used ‘STRING Experimental PPI’ option for the knowledge network (which corresponds to experimentally verified protein-protein interaction edges from the STRING database [14]) and the amount of network smoothing was set to the default 50%. Only pathways with ‘difference score’ larger than 0.5, which correspond to those that have a query score higher than the background score, were considered associated with the input query gene set. For enrichment analysis of the set of TFs, the input file to KnowEnG’s GSC pipeline was designed such that the universe (i.e. population) would be equal to the set of all TFs present in our study. For enrichment analysis of a TF and its target genes, the universe was set to be the set of all genes and TFs present in our study.

## Authors’ Contributions

AE and SR conceived the study. CS performed the analyses. All authors contributed to the drafting of the manuscript and critical discussion of the results. All authors read and approved the final manuscript.

## Acknowledgements

This work was supported by Natural Sciences and Engineering Research Council of Canada (NSERC) grant RGPIN-2019-04460 (AE) and McGill Initiative in Computational Medicine (MiCM) (AE and SR).

## Titles of Supplementary Files

**Supplementary Figure S1:** The histogram of z-score normalized gene expression changes of LINCS L1000 landmark genes in A549 cells as a result of knockdown of kinases implicated in the SvM analysis. Each histogram corresponds to the knockdown of one kinase. Vertical lines depict the 15th and 85th percentiles and red stars show the TFs and target genes in the SvM network that are positively or negatively influenced by the experiment.

**Supplementary Figure S2:** The effect of kinase knockdowns on genes and TFs shared between the SvM TRN and L1000 landmark genes. An upward arrow shows that the expression of the gene in A549 cells increased and the change in the expression was among the top 15% of all landmark genes. A downward arrow shows that the expression of the gene in A549 cells decreased and the change in the expression was among the bottom 15% of all landmark genes.

**Supplementary Figure S3:** The effect of kinase knockdowns on genes and TFs shared between the SvOV TRN and L1000 landmark genes. An upward arrow shows that the expression of the gene in A549 cells increased and the change in the expression was among the top 15% of all landmark genes. A downward arrow shows that the expression of the gene in A549 cells decreased and the change in the expression was among the bottom 15% of all landmark genes.

**Supplementary Table S1**: Top 500 differentially expressed genes and their corresponding p-values. First tab corresponds to genes differentially expressed between SARS-CoV-2 infected cells and mock-treated cells. The second tab corresponds to genes differentially expressed between SARS-CoV-2 infected cells and cells infected by other respiratory viruses.

**Supplementary Table S2**: The SvM and SvOV TRNs constructed using InPheRNo. Column headers correspond to TFs and row names correspond to target genes. Each entry in the matrix reflects the score assigned to each potential TF-gene edge. Only edges with score > 0.5 were considered for the follow-up analysis.

**Supplementary Table S3**: The full list of TFs ranked based on the number of COVID-19-relevant target genes identified by InPheRNo in the SvM and SvOV networks.

**Supplementary Table S4**: Results of standard and network-guided pathway enrichment analysis using KnowEnG’s GSC computational pipeline. The results of the first two tabs are obtained by selecting the top 8 TFs from the SvM network as the query gene set, while the results of the last two tabs are obtained by selecting the top 21 TFs from the SvOV network as the query gene set.

**Supplementary Table S5**: Network-guided pathway enrichment analysis corresponding to TFs and target genes identified by InPheRNo in the SvM network. The results of each tab are obtained by using an implicated TF and its identified target genes as a query gene-set.

**Supplementary Table S6**: Network-guided pathway enrichment analysis corresponding to TFs and target genes identified by InPheRNo in the SvOV network. The results of each tab are obtained by using an implicated TF and its identified target genes as a query gene-set.

**Supplementary Table S7**: The full ranked list of kinases identified by foRWaRD for the SvM and SvOV networks.

**Supplementary Table S8**: The indirect interactions of the implicated kinases and TFs in the SvM and SvOV networks corresponding to Fig. 3. The kinase-substrate relationships are directly extracted from the aggregated KSI. The substrate-TF interactions correspond to edges in the HumanNet Integrated network.

**Supplementary Table S9**: The effect of shRNA knockdown of implicated kinases on the TFs and target genes identified by InPheRNo in A549 cells. Each tab includes the z-score normalized changes in the expression of the genes as well as the percentile of that change among all L1000 landmark genes.

