## Supplementary figures and images for "Identification of COVID-19-relevant transcriptional regulatory networks and associated kinases as potential therapeutic targets"

### Supplementary Figure S1

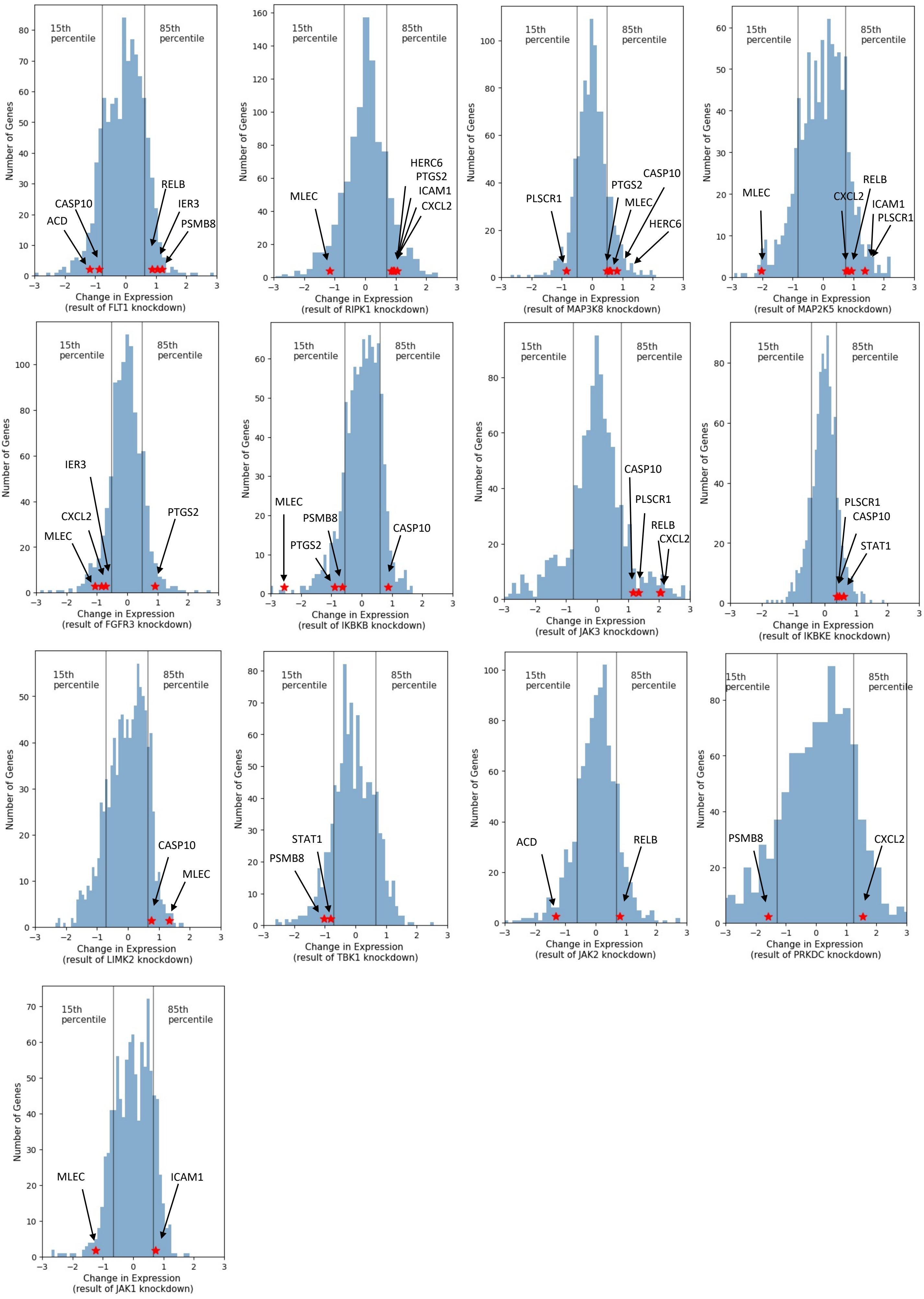

### Supplementary Figure S2

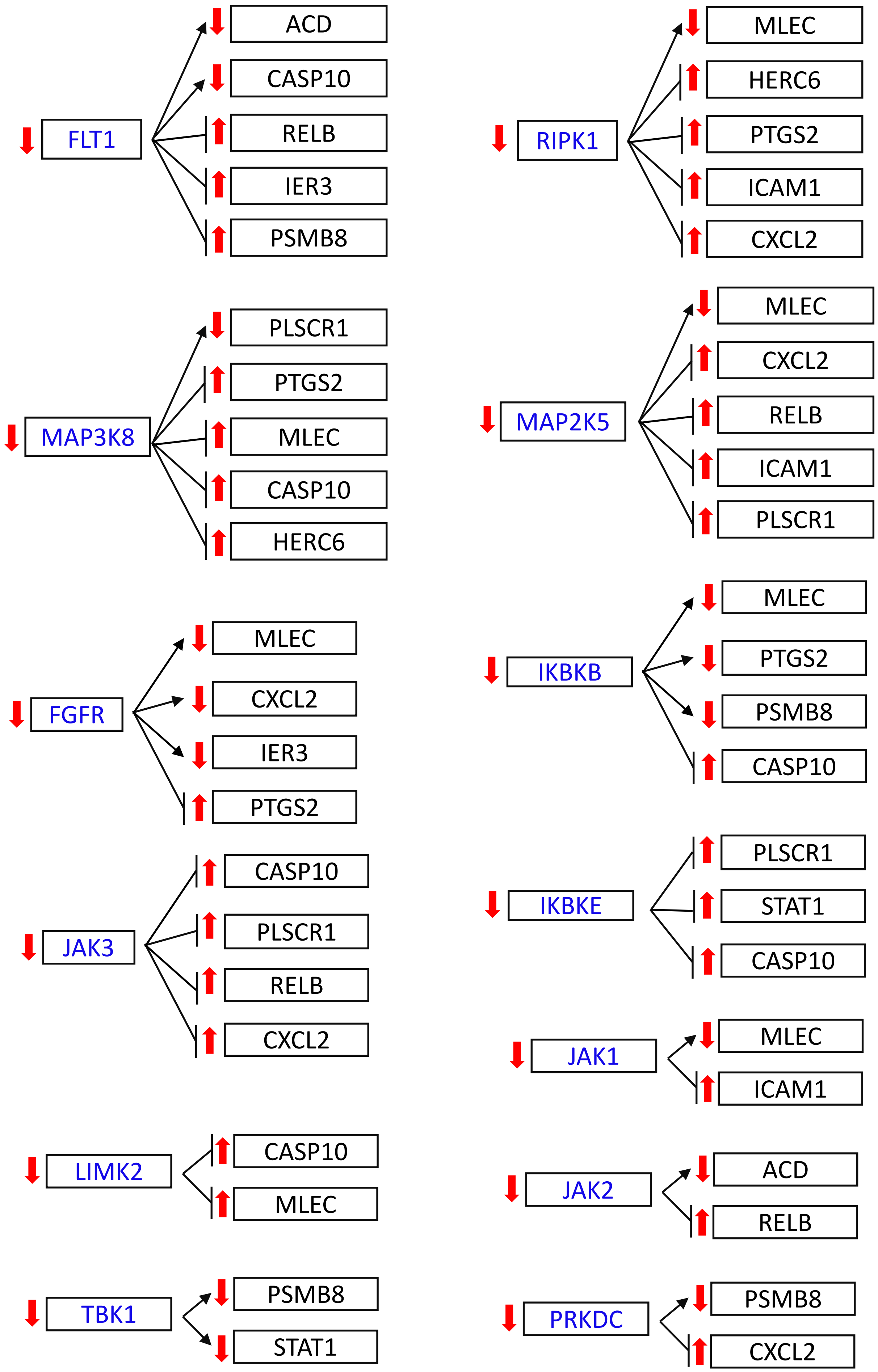

### Supplementary Figure S3

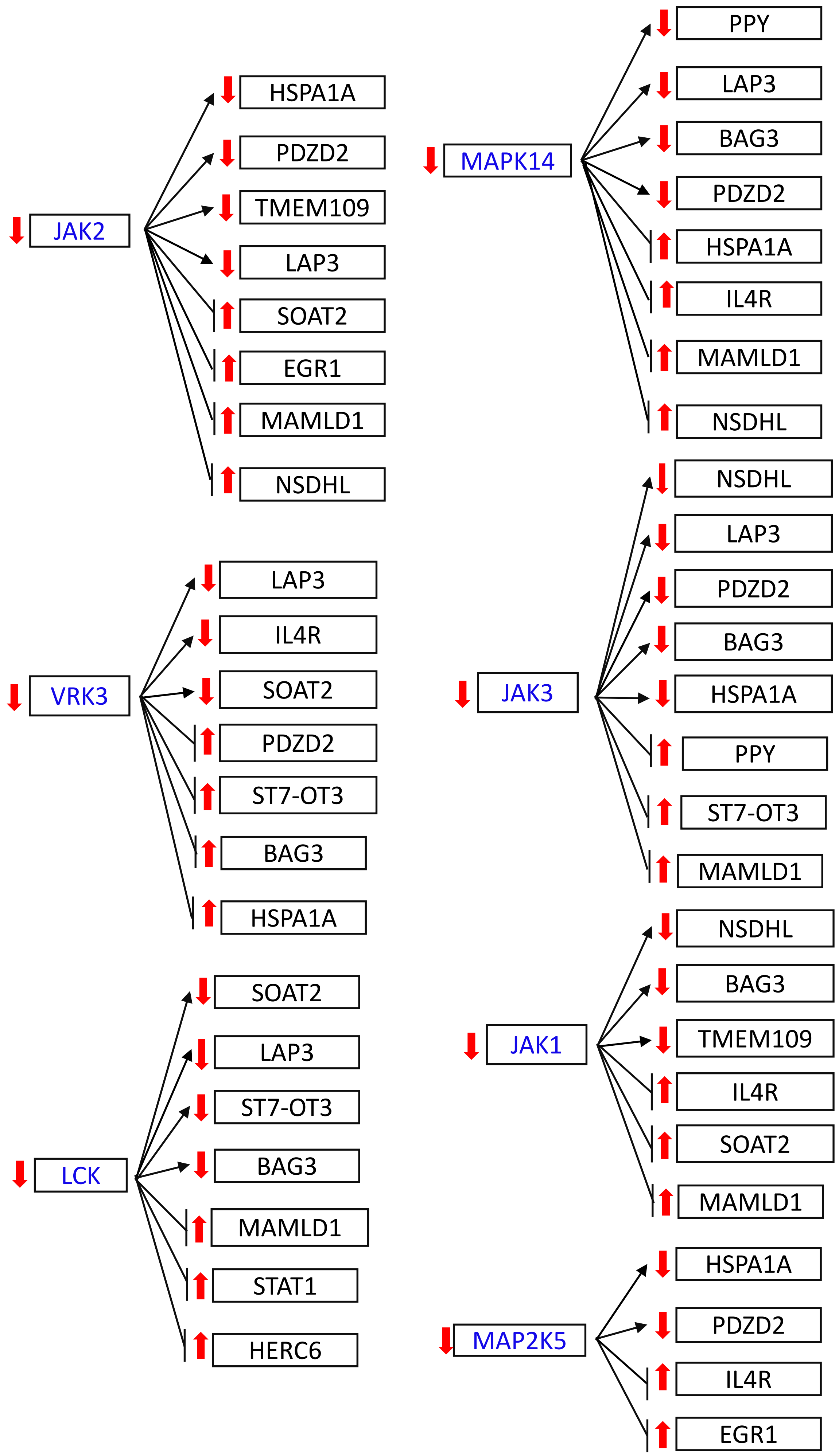
